# Decoding the “bouba-kiki” effect in early visual cortex

**DOI:** 10.64898/2026.01.17.700088

**Authors:** Stefano Ioannucci, Carole Jordan, Anne-Sophie Carnet, Carolyn McGettigan, Petra Vetter

## Abstract

The “bouba-kiki” effect, or sound symbolism, is the non-arbitrary mapping between speech sounds and visual shapes, in which words like “bouba” are consistently associated with round shapes, and “kiki” with spiky shapes. Despite robust behavioral evidence for sound symbolic associations, it is still unclear whether they are a linguistic epiphenomenon or manifest themselves as distinct sensory representations in the human brain. Using fMRI and multivariate pattern analyses, we show that, in the absence of visual input, round and spiky sound symbolic words elicit distinct neural representations, not only in auditory and speech processing regions, but also in parietal and even early visual cortex including V1. These neural representations correlate with behavioural shape judgements of sound-symbolic words, as revealed by representational similarity analyses. Our findings suggest a potential functional mechanism for sound symbolism: phonological features of sound symbolic words are represented in auditory and speech processing regions and linked to cross-modal shape representations in parietal and early visual regions, which in turn lead to sound symbolic perceptual judgements. Thus, our results provide evidence for a neural mechanism underlying the “bouba-kiki” effect in the human brain.

## Introduction

The “bouba-kiki” effect, also referred to as sound symbolism, is a century-old phenomenon where certain speech sounds are consistently associated with particular visual features (Köhler, 1929; Sapir, 1929; Ramachandran & Hubbard, 2001). For example, when participants are asked to match meaningless words to visual shapes, they consistently map words like “bouba” to round shapes, and words like “kiki” to spiky shapes. Beyond shape, similar mappings, or cross-modal correspondences, exist between speech sounds and size, brightness, color, and other visual features (Spence, 2011; Lockwood & Dingemanse, 2015; Sidhu & Pexman, 2018). Importantly, cross-modal sound symbolic associations have been observed across developmental stages (Maurer et al., 2006; Ozturk et al., 2013) and across diverse populations, cultures and writing systems (Bremner et al., 2013; Ćwiek et al., 2021; Thompson et al., 2021), suggesting a potentially universal phenomenon independent of a written language system and the ability to read.

Despite a long tradition of robust behavioural evidence for the “bouba-kiki” effect, it is still unclear whether this effect is a linguistic epiphenomenon or manifests as a true cross-modal effect in the human brain with distinct neural representations in sensory cortices. Neural evidence for sound symbolism is still relatively sparse and mixed. Associating sound-symbolic words with abstract visual properties preferentially activates parietal cortex compared to non-sound-symbolic words (Revill et al., 2014). When sound-symbolic words match simultaneously presented visual shapes and are compared to non-matching audiovisual stimuli, occipital EEG activity is enhanced (Kovic et al., 2010) and auditory, prefrontal and fusiform/parahippocampal fMRI activity is elevated (Peiffer-Smadja & Cohen, 2019; Barany et al., 2023).

However, it remains unknown whether sound symbolic speech sounds alone, without visual stimulation, elicit distinct representations in visual cortices solely due to their strong cross-modal associations with visual shape properties. If this was the case, then the “bouba-kiki” effect would manifest itself with distinguishable representations for different sound symbolic words in visual cortices even in the absence of visual input.

Recent neuroscientific investigations have expanded our understanding of the role of human visual cortex beyond its established visual functions, particularly with respect to its representation of auditory features (Brang et al., 2015; Murray et al., 2016; Petro et al., 2017). For example, natural sound categories can be decoded from early visual cortex activity in blindfolded sighted (Vetter et al., 2014; Pollicina et al., 2025), aphantasic (Montabes de la Cruz et al., 2024) and congenitally blind (Vetter et al., 2020) individuals.

This evidence challenges traditional views of sensory cortices as purely modality-specific and reveals how the different senses, perception and cognition may be intertwined by more complex integrative or predictive processes (Anderson, 2010; Friston, 2010; Petro et al., 2014; Vetter et al., 2014; Vetter & Newen, 2014; Newen & Vetter, 2017). Sound symbolism offers an ideal setting to investigate whether top-down predictions stemming from audition, such as implicit shape associations in speech sounds, can influence neural activity in visual cortex. Given the robustness of the “bouba-kiki” effect, we investigated whether sound-symbolic words would elicit distinguishable activity patterns in early visual cortex regions, depending on their shape association (round versus spiky), even in the absence of visual stimulation.

We conducted two fMRI experiments in which blindfolded adult participants either judged the shape of sound-symbolic words or their meaningfulness. Moving beyond traditional paradigms that employed only purely round or purely spiky words, we included mixed-category words (e.g. “baki”) to investigate whether sound-symbolic word representations are driven by phonology or by visual shape associations. Using multivariate pattern analysis (MVPA; Haxby et al., 2014), we tested whether neural activity patterns in early visual cortex distinguish between different sound-symbolic word categories. We complemented a region-of-interest (ROI) approach with a whole-brain searchlight analysis (Etzel et al., 2013) to identify relevant clusters in brain areas beyond our a-priori anatomical regions of interest.

We hypothesized that if the “bouba-kiki” effect is a true effect of differential sensory representations, then round and spiky words should be successfully decodable from fMRI activity patterns in both early visual areas (V1, V2, V3) and auditory cortex (A1 [TE1.0], A2 [TE1.1], A3 [TE1.2]), also in the absence of visual input. We also assessed whether these shape categories would be decodable in visual shape-sensitive lateral occipital complex (LOC), speech-sensitive Broca’s area (BRC) and in control regions which could explain motor confounds (post-central gyrus; PCG) or visual imagery of the written form of the pseudowords (visual word form area; VWFA). For the whole-brain searchlight analysis, we expected to identify discriminative clusters in higher visuo-spatial or cross-modal regions previously implicated in sound-symbolic associations.

Additionally, we predicted a correspondence between perceptual and neural representations of sound-symbolic categories. To test this relationship, we employed representational similarity analysis (RSA; Kriegeskorte et al., 2008) to quantify representational distances between categories in behavioral shape judgments and correlate these with distances between neural representations of the same categories.

## Results

### Behavioral judgment of sound-symbolic shape

In the main experiment, 24 blindfolded healthy adult French speakers judged the shape of sound symbolic words while laying inside a 3T MRI scanner. To examine the task-specificity of sound symbolic representations, an additional cohort of 22 participants judged the meaningfulness of identical stimuli in a second experiment. The speech stimuli were validated in behavioural pilot experiments on a separate sample (N = 34) and consisted of 24 unique bisyllabic spoken pseudowords (without meaning in French) across 4 shape categories and 12 meaningful words (in French) composed of similar syllables (see Table 1 in Methods). The shape categories were: 1) round words (e.g. “doba”) composed entirely of syllables typically associated with roundness (Nielsen & Rendall, 2013), 2) spiky words (e.g. “kipe”) composed entirely of syllables typically associated with spikiness. 3) mixed round words (e.g. “bopi”) containing one round-associated syllable and one spiky-associated syllable and that were judged as mostly round in the behavioural pilot experiment, and 4) mixed spiky words (e.g. “tiba”) containing one spiky-associated syllable and one round-associated syllable that were judged as mostly spiky in the pilot. The 12 meaningful words were composed of the same type of syllables with both pure and mixed shape ratings (e.g. “dodo”). Words were presented in pseudo-randomised order via headphones. In the main experiment, participants rated the shape of the words on a scale from 1-4 (most round to most spiky, or vice versa, counterbalanced across participants). In the second experiment, participants indicated whether the word had a meaning in French or not. In both experiments, participants were blindfolded, instructed to keep their eyes closed at all times and room lights were switched off, ensuring as little visual input as possible.

**Table 1:**
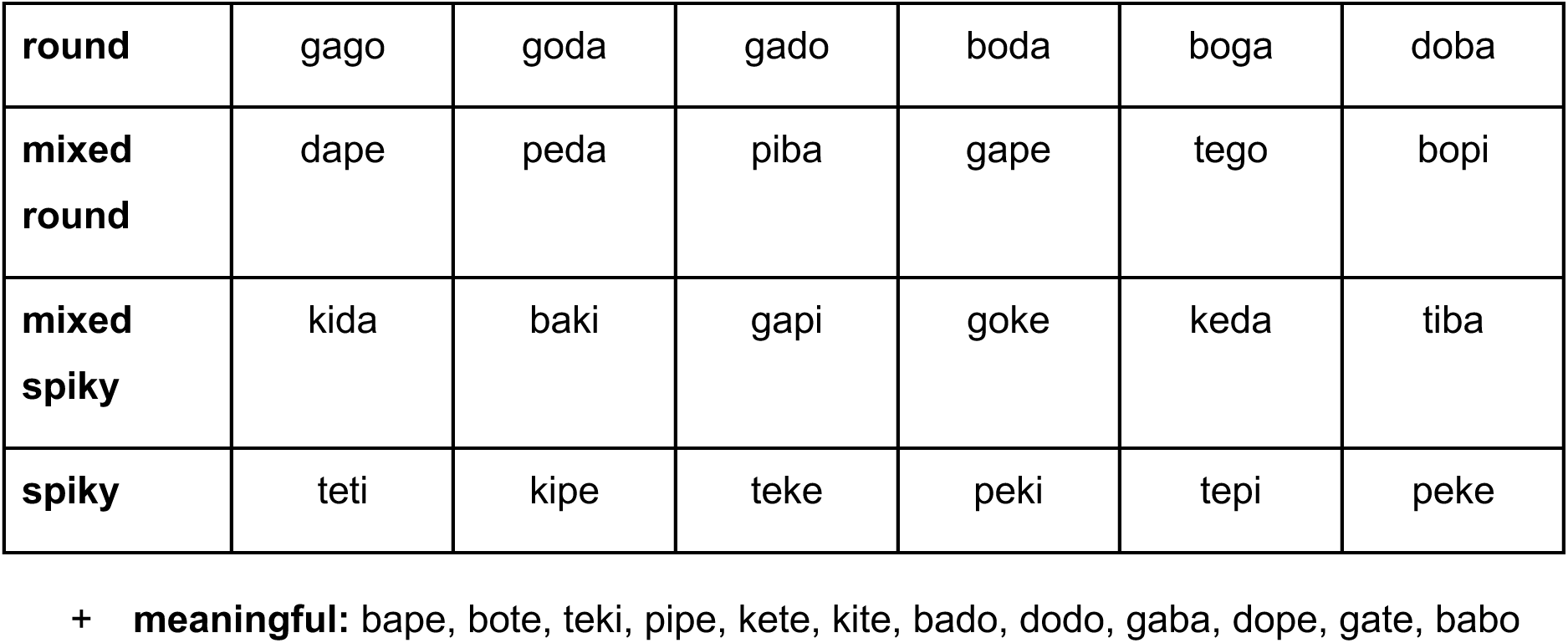
Sound-symbolic word stimuli and their categorization by shape. Words were selected based on results of behavioural pilot experiments in which words in the round category were consistently judged as maximally round, in the mixed round category as mostly round, in the mixed spiky category as mostly spiky, and in the spiky category as maximally spiky. 12 meaningful words (in French) were also included in the stimulus sample.

Behavioural shape judgements of spoken words inside the scanner demonstrated a clear “bouba-kiki” effect, with sound-symbolic words being clearly associated with their implied shape (F_(3)_ = 142, *p* < 0.001, all post-hoc comparisons *p_corr_* < 0.001; **Figure 1**), with a highly consistent gradient from round to spiky judgements along round, mixed-round, mixed-spiky and spiky categories. The mean difference in shape ratings between purely spiky and purely round words was substantial (Δ = 1.48, Cohen’s d = 2.85), approximately three times larger than the differences between adjacent categories on the gradient (spiky vs. mixed spiky: Δ = 0.41, d = 1.4; mixed spiky vs. mixed round: Δ = 0.49, d = 1.6; mixed round vs. round: Δ = 0.58, d = 1.5). This behavioural response pattern demonstrated highly robust sound symbolic associations and a graded influence of phonological features on visual shape associations.

**Figure 1:**
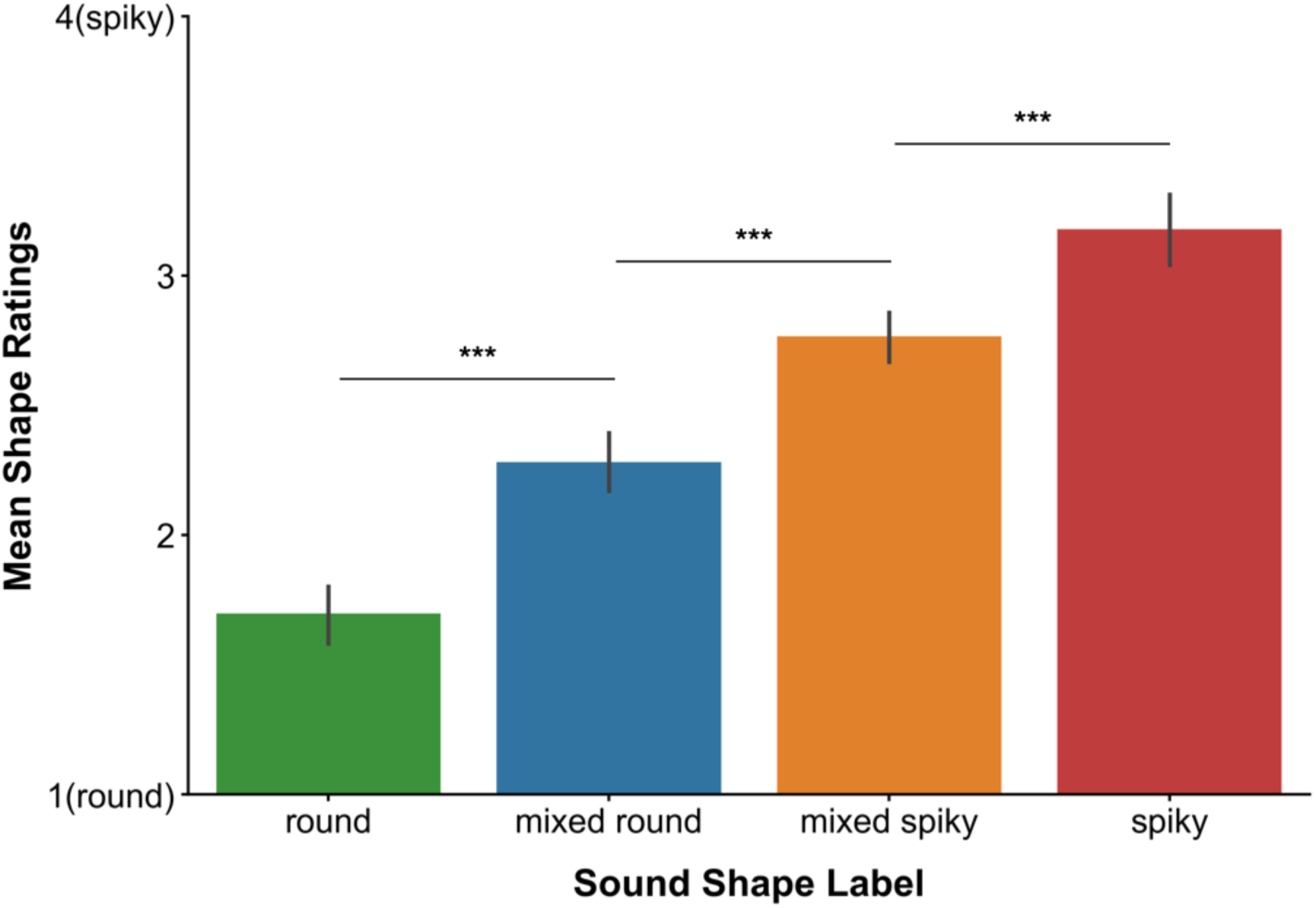
Behavioral mean shape ratings (y axis; 1 = round, 4 = spiky) of spoken sound-symbolic words by shape category (x axis; colors)

### Decoding of sound-symbolic words

For each sound-symbolic word that blindfolded participants rated, single-trial brain activity patterns were estimated (Prince et al., 2022) and then fed into a linear classifier for 4 different MVPA classification pairs with leave-one-run-out cross-validation: round vs spiky, mixed round vs mixed spiky, round vs mixed round and spiky vs mixed spiky. These classifications were carried out on normalised fMRI BOLD signals within anatomically defined ROIs (Dehaene et al., 2002; Wang et al., 2015; Benson & Winawer, 2018; Amunts et al., 2020), encompassing early visual cortex (EVC: V1, V2, V3), lateral occipital regions (LO1, LO2), primary and secondary auditory cortices (AC: A1, A2, A3), speech processing Broca’s area, and control regions (PCG, VWFA).

When participants explicitly judged the implied shape of sound-symbolic words (Experiment 1), group-level classification analyses revealed distinct neural representations for sound-symbolic spoken words in selected brain regions. Classification between round and spiky pseudowords (**Figure 2**) succeeded significantly above chance in early visual cortex, particularly in V1 (Z = 3.7, *p_corr_* = 0.007, d = 0.7), in auditory cortex, particularly in A3 (Z = 6, *p_corr_* < 0.001, d = 1.3) and in Broca’s area (Z = 2.5, *p_corr_* = 0.025, d = 0.6). Thus, neural activity patterns in early visual cortex, as well as in primary auditory cortex and Broca’s area, contained distinguishable representations for sound-symbolic spoken words that were maximally distinct in terms of implied shape (purely round versus purely spiky). Importantly, in none of these ROIs was successful classification driven by differences in mean univariate BOLD activity between round and spiky words (**Fig. S1**; all *p_corr_* > 0.18). This means that the MVPA classifier picked up on the distinct patterns of distributed small activity differences in these regions rather than mean univariate activity levels that could have been driven by unspecific factors like differential attention, arousal or imagery. Instead, the classification results indicate *distinct neural representations* of round and spiky words in V1, auditory cortex and Broca’s area.

**Figure 2:**
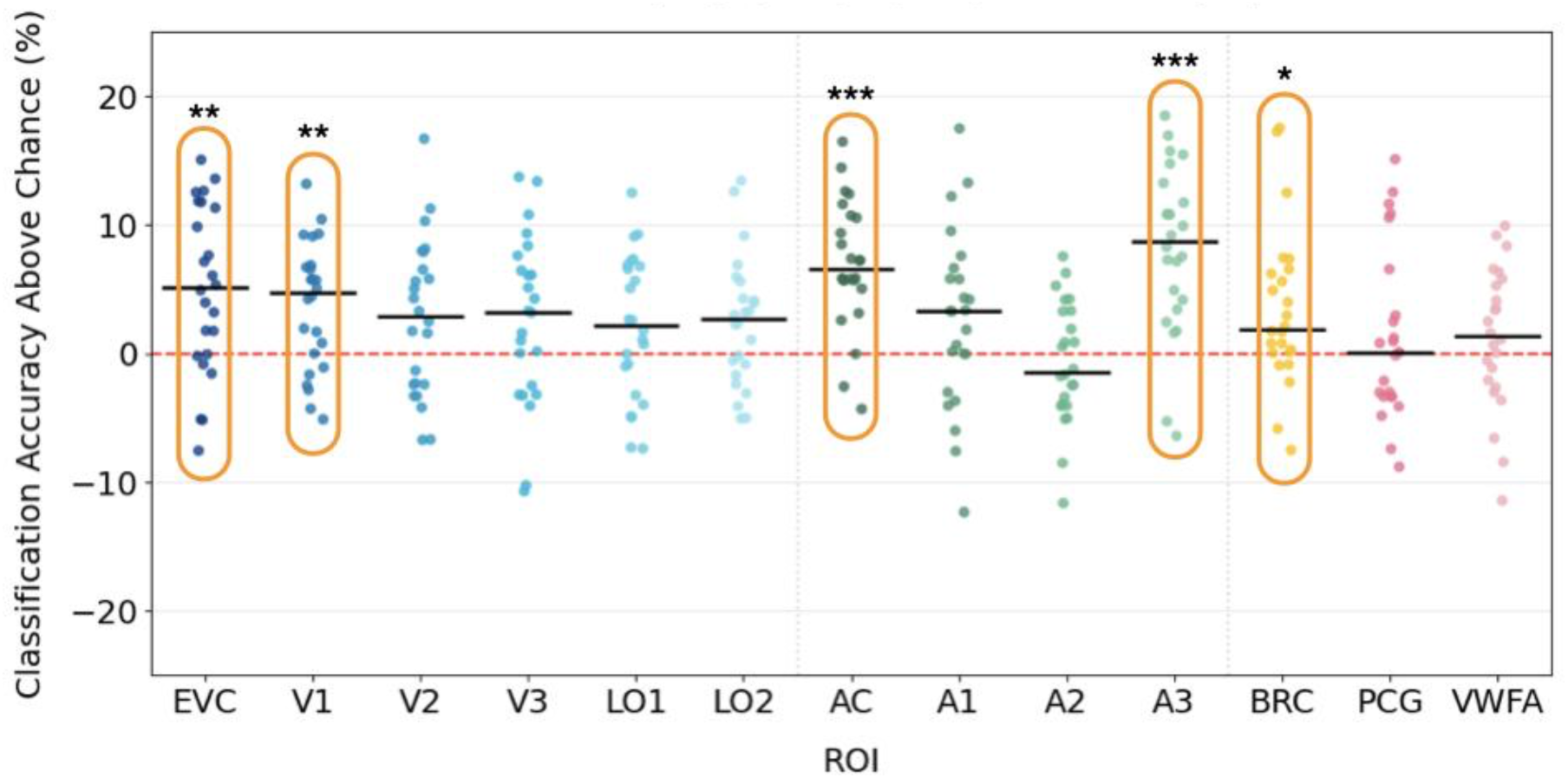
Chance-subtracted MVPA classification accuracies (y axis) for each participant (dots) and ROI (x-axis) for the classification between round and spiky spoken words during shape judgment (Experiment 1). The red dashed line represents chance level, horizontal black bars represent the group-wise medians and orange contours highlight significant ROIs (*p < .05; ** p < .01; *** p < .001).

Classification of the mixed word categories with any of the other categories was not successful in any of our ROIs (mixed spiky vs spiky: all *p_corr_* > 0.22, mixed round vs round: all *p_corr_* > 0.08 mixed round vs mixed spiky: all *p_corr_* > 0.28), suggesting that neural representations of round and spiky words require clear differences in either shape associations or phonology to be discernible.

We also did not find significant above-chance classifications in any of the control regions VWFA or PCG (see Supplementary Table S1) suggesting a highly specific effect in early visual and auditory cortices and Broca’s area. Unsuccessful classification in the VWFA makes it unlikely that visual imagery of the written word form had strongly driven successful classification in V1. Similarly, unsuccessful classification in PCG makes it unlikely that differential button responses to round and spiky words drove classification in other brain areas. Furthermore, we did not find significant correlation in subject-wise classification accuracies between V1 and A3 in the round versus spiky classification (R_(22)_ = 0.01, *p* = 0.95), suggesting that V1 and A3 may have complementary roles in representing sound-symbolic words, each exploiting different auditory and visual features to distinguish the sounds and their associated shape.

To test task-specificity of sound-symbolic word representations, we conducted a second fMRI experiment in which a separate cohort of 22 blindfolded participants listened to identical stimuli, but instead of performing a shape judgement task, they judged whether the words had a meaning in French or not. Here, classification of round versus spiky words succeeded only in auditory cortex (A1: Z = 3.7, *p_corr_* < 0.001, d = 1.2; A3: Z = 2.9, *p_corr_* = 0.016, d = 0.72; **Figure 3**) consistent with the processing of inherent acoustic-phonetic differences between round and spiky word categories that persist regardless of task demands. Again, successful classification was not driven by differences in univariate activity levels in the relevant ROIs (all *p_corr_* > 0.7; **Figure S2**). As in the first experiment, none of the other shape category classifications succeeded (mixed spiky vs spiky: all *p_corr_* > 0.11, mixed round vs round: all *p_corr_* > 0.08, mixed round vs mixed spiky: all *p_corr_* > 0.43; see Supplementary Table S2). The results of the second experiment suggest that distinct representations of sound-symbolic round and spiky words do not emerge in early visual cortex when shape associations are task-irrelevant.

**Figure 3:**
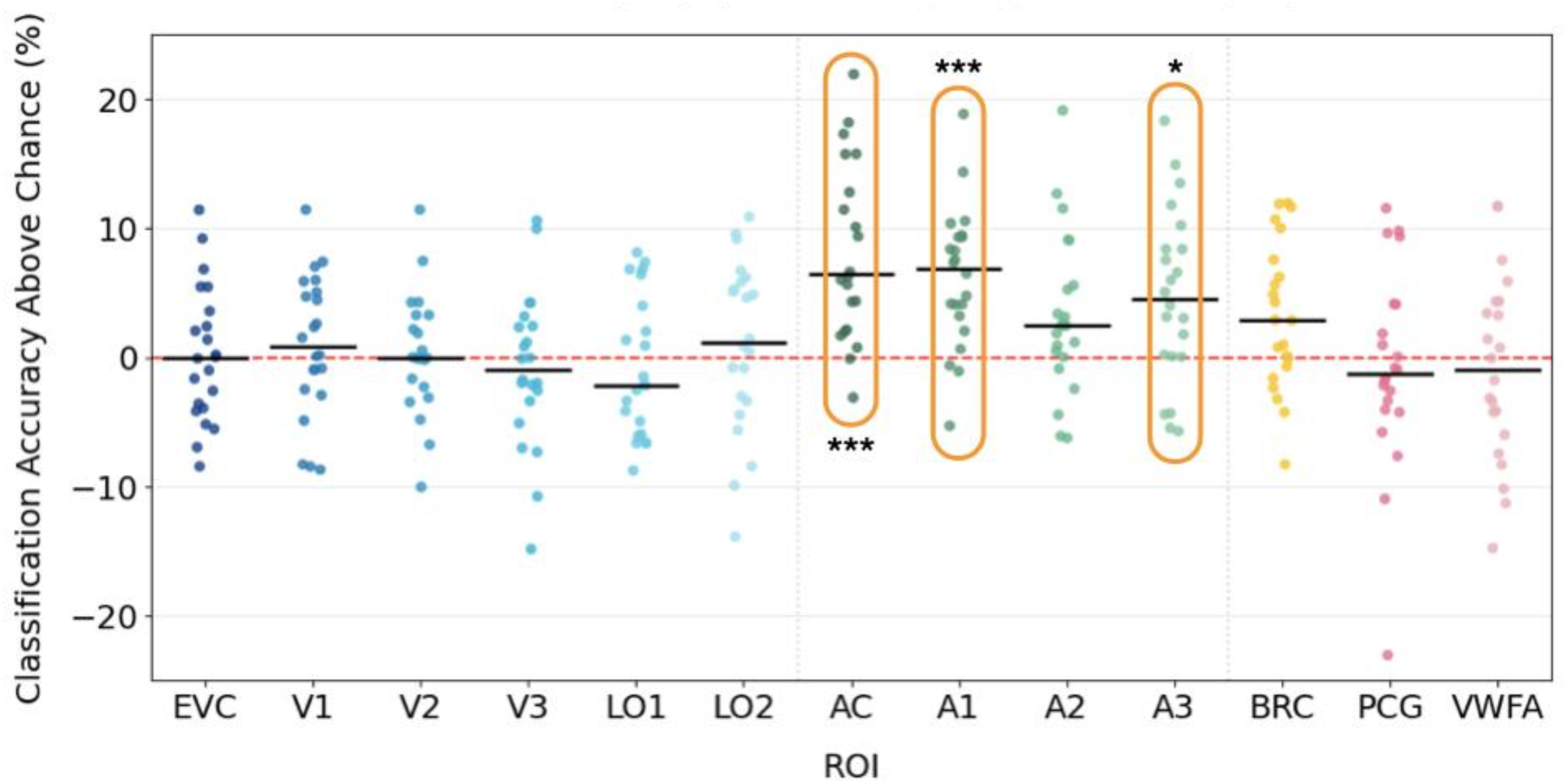
Chance-subtracted MVPA classification accuracies (y axis) for each participant (dots) and ROI (x-axis) for the classification between round and spiky spoken words during meaningfulness judgments (Experiment 2). The red dashed line represents chance level, horizontal black bars represent the group-wise medians and orange contours highlight significant ROIs (*p < .05; ** p < .01; *** p < .001).

### Whole-brain Searchlight Results

We also conducted a whole-brain voxel-wise searchlight decoding analysis to localize clusters that were most spatially consistent across participants in displaying above chance classification accuracy distinguishing round and spiky words (Exp.1). Two significant clusters were identified and located, via Julich cytoarchitectonic atlas (Amunts et al., 2020), in the right intraparietal sulcus (right IPS; [K_z_ = 1.73, *p_corr_* = 0.019, K = 129]) and the left superior temporal gyrus (left STG; [K_z_ = 2.31, *p_corr_* = 0.005, K = 225], **Figure 4)**. The cluster in left superior temporal gyrus overlapped with our auditory cortex ROI (particularly A1 and A3, Figure S3). Successful classification of round versus spiky words in left STG and right IPS is consistent with their involvement in complex auditory (Hickok & Poeppel, 2007) and visuospatial processing (Culham & Kanwisher, 2001; Xu & Jeong, 2016), respectively.

**Figure 4:**
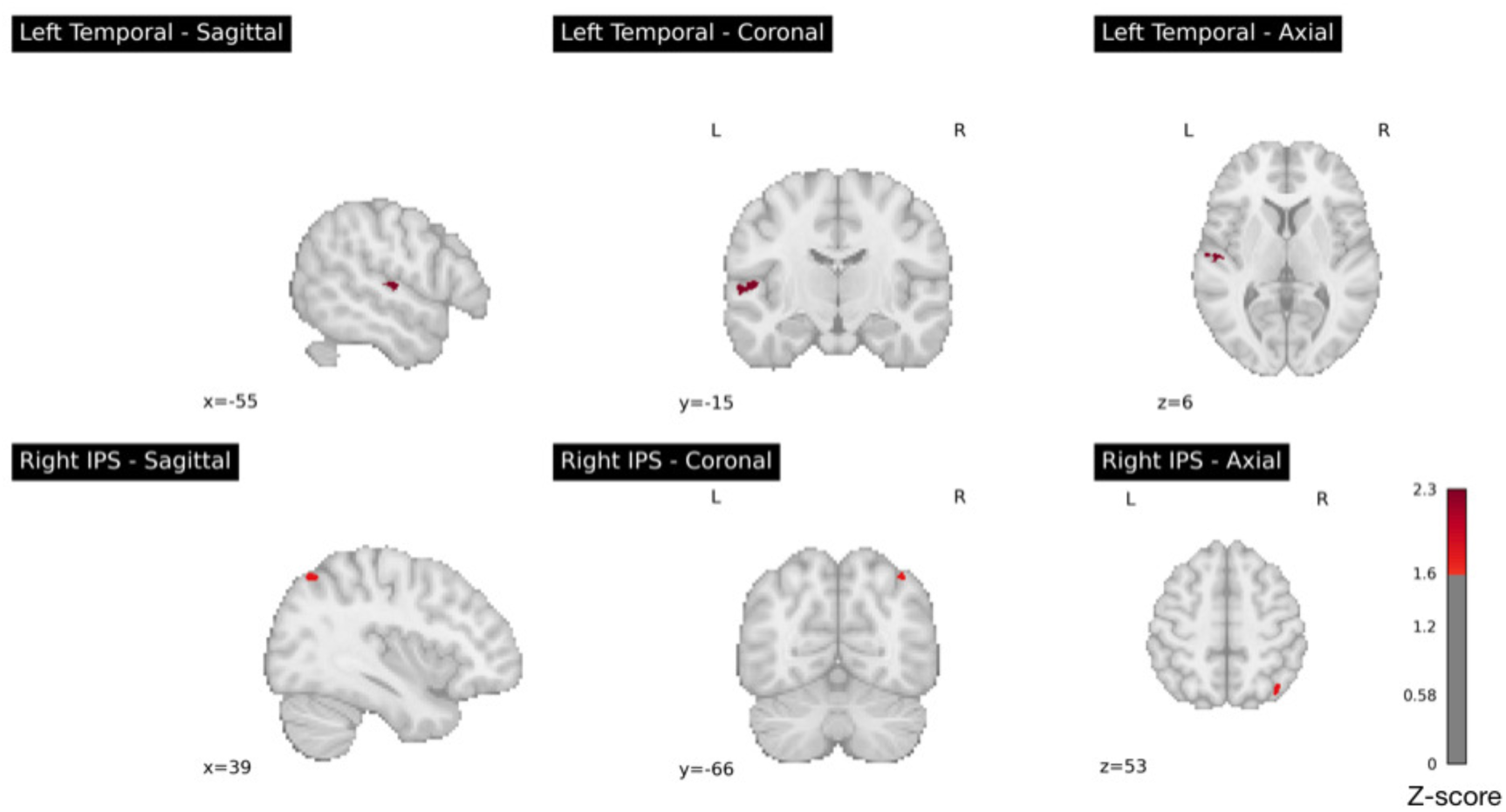
Group-level Z-maps of the whole-brain searchlight classification results (Experiment 1). Round versus spiky words could be classified significantly above chance in the shape judgement experiment in left Superior Temporal Gyrus (top row) and right Intraparietal Sulcus (bottom row). Significant clusters (red) indicate group-level above chance classification *p_corr_* < .05.

Notably, exploration with a more liberal statistical threshold revealed only the contralateral homologues of these regions (right STG; *p_corr_* = 0.12, left IPS; *p_corr_* = 0.09), making the above effects unlikely to be strongly lateralised.

### Representational geometry between behavior and neural signal

To quantify how similarly all shape categories were represented across neural activity and behaviour, we calculated the representational dissimilarity matrices for the behavioral judgments and for the neural representations within the relevant ROIs, i.e. those with above-chance classification in the MVPA and searchlight analyses (Exp. 1). These matrices were then correlated for each participant, and their correlation coefficients (Kendall’s τ) were tested via sign-rank tests, corrected for multiple comparisons across ROIs. This analysis revealed that, at the group level, the representational geometry was positively correlated between the neural activity patterns and behavior in V1 (Z = 3.1, *p_corr_* = 0.048, d = 0.7) and left STG (Z = 3.2, *p_corr_* = 0.048, d = 0.7; **Figure 5**).

**Figure 5:**
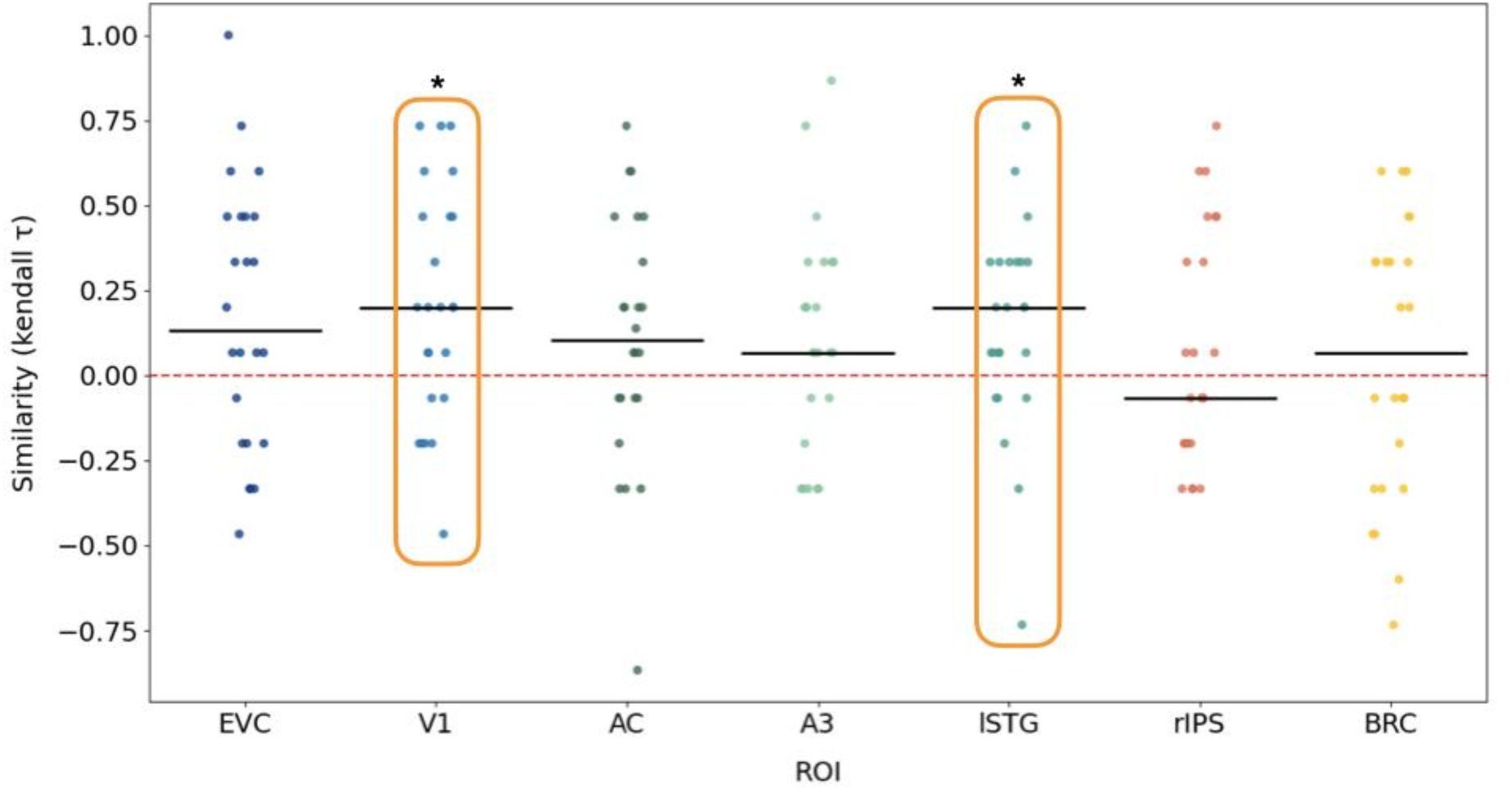
Results of the RSA analysis. Kendall’s tau correlation coefficients between the representational geometry of shape categories between neural activity and behavioral shape judgment (y-axis) for each participant (dots) and ROI (x-axis). The red dashed line represents no correlation, horizontal black bars represent the group-wise medians, and orange contours highlight ROIs with significant correlations (*p < .05).

### Decoding meaningfulness of words

We also analysed whether meaningless versus meaningful words elicited distinguishable representations in both the first and second experiment. In the second experiment, in which participants judged whether the presented word had a meaning in French or not, behavioral performance confirmed successful task engagement, with participants reliably rejecting meaningless words compared to accepting meaningful ones (60 ± 15 % vs 10 ± 4 %, chance = 20%, T_(21)_ = -13, *p* < 0.001). MVPA classification of meaningful versus meaningless words succeeded exclusively in Broca’s area (Z = 3.7, *p_corr_* < 0.001, d = 1.2; Figure S4) when task-relevant in Experiment 2, while the same classification failed in all other ROIs, as well as in all ROIs during shape judgement in Experiment 1 (all *p_corr_* > 0.27). This suggests that representations of word meaning of these specific pseudowords only emerged in Broca’s area when task-relevant in Experiment 2.

## Discussion

The “bouba-kiki” effect has been studied for almost a century in psychology (Köhler, 1929; Sapir, 1929; Newman, 1933; Ramachandran & Hubbard, 2001), and while it is behaviourally very well established (Spence, 2011; Lockwood & Dingemanse, 2015; Sidhu & Pexman, 2018), evidence for its sensory representations and neural mechanism in the human brain is still sparse. Using MVPA brain decoding techniques, we demonstrate here that, during shape judgements, early visual cortex, including primary visual cortex V1, represents “round” and “spiky” sound-symbolic words in a distinctive manner even in the absence of visual input. Round and spiky words also elicit distinguishable representations in right intraparietal cortex, auditory regions and Broca’s area. Thus, the “bouba-kiki” effect manifests itself with distinct cross-modal neural representations in the human brain and is not just a linguistic epiphenomenon.

While previous studies demonstrated enhanced neural activity in some cross-modal and sensory cortices in response to sound-symbolic words when presented concurrently with visual stimuli (Revill et al., 2014; Peiffer-Smadja & Cohen, 2019), our approach distinguishes different neural representations of sound-symbolic shape categories (“round” versus “spiky”) in several sensory and cognitive brain regions. When shape properties are task-relevant, different sound-symbolic shape associations of spoken words are sufficiently robust to elicit distinct neural activity patterns in V1 even without any actual visual stimulation. This shows

1. that sound-symbolic associations do not just happen on a cognitive or linguistic level, but are distinctively represented in primarily sensory regions like V1 and primary auditory cortex;
2. that V1 is capable to maintain differential representations of meaningless speech sounds if they are associated with visual shape categories. These findings provide evidence for a potential neural mechanism on how sound symbolism comes about, namely by co-representing sound-symbolic shape categories (“round” and “spiky”) in both primary visual and primary auditory cortex, as well as in right intraparietal cortex and Broca’s area.

The second experiment, in which participants judged meaningfulness instead of shape of the words, revealed that sound symbolic representations are not automatically elicited in visual cortex, but instead flexibly elicited depending on task demands. When participants judged the meaning of word stimuli, visual cortex activity no longer carried sufficient information to distinguish between round and spiky words. In contrast, auditory cortex maintained distinct representations regardless of task demands. Such dissociation suggests that auditory cortex likely distinguishes primarily the acoustic-phonetic differences between sound-symbolic word categories, while V1 represents cross-modally associated shape only when behaviorally relevant. Furthermore, the pattern of representations in Broca’s area was also modulated by

task demands. During shape judgment, Broca’s area showed distinguishable representations for round versus spiky words, suggesting engagement in sound-symbolic speech processing, in line with categorical speech processing (Lee et al., 2012). However, during judgment of meaningfulness of identical stimuli, Broca’s area instead distinguished meaningful versus meaningless words, in accordance with previous evidence (Abrams et al., 2013), while losing the capacity to distinguish shape categories. This complementary dissociation provides compelling evidence that both early visual cortex and Broca’s area flexibly adapt their representational content to match current cognitive demands.

Our findings reveal that sound symbolism relies on flexible, cognition-gated recruitment of both sensory and speech processing regions rather than fixed cross-modal mechanisms. The capacity for the same neural populations to represent different stimulus dimensions based on task relevance mirrors accumulating evidence for the impact of task on vision and visual cortex (Kay et al., 2023). Our results exemplify this remarkable flexibility of sensory regions to adapt to cognitive task demands in the context of cross-modal perception (see also Vetter et al., 2024).

Our behavioral results show that participants consistently associated sound-symbolic words (e.g. “boda”, “teke”) with their implied round or spiky visual shapes, despite having no prior exposure to these stimuli and being naive to the “bouba-kiki” effect. This corroborates once more the robustness of the intrinsic sound-shape mappings underlying sound symbolism (Spence, 2011; Lockwood & Dingemanse, 2015; Sidhu & Pexman, 2018). Furthermore, participants’ shape judgments were consistent also for ‘mixed’ stimuli containing a round and spiky syllable (e.g. “dape”, “kida”), with the mixed-round stimuli being judged as more round than the mixed-spiky category, confirming the results of the behavioural pilot experiment.

The systematic gradient of shape judgements across stimulus categories suggests that sound symbolism operates along a perceptual continuum rather than along discrete category boundaries, with a graded mapping of acoustic features to visual properties. In line with this graded mapping, mixed-category words did not elicit distinguishable representations in neither visual nor auditory cortices, neither when contrasted with the other mixed category nor with any of the pure “round” or “spiky” categories. This was likely due to too similar representations, both on a phonological/acoustic level and on the level of visual shape associations, not allowing successful distinction of neural activity patterns by the MVPA classifier.

While “pure” round and spiky sound-symbolic words could be decoded significantly above chance in early visual and auditory cortices, they could not be classified in the visual word form area (VWFA). This makes it unlikely that participants’ potential visualization of the written word forms strongly drove successful decoding in V1. Additionally, the fact that classification accuracies in visual and auditory cortices did not correlate may suggest independent representations in these sensory regions, indicating that visual areas may preferentially distinguish implied shape and auditory regions may preferentially distinguish acoustic features. In fact, successful decoding of sound-symbolic word categories in auditory cortices, independent of task, suggests that decoding in auditory cortex was likely driven by acoustic and phonological differences between word categories.

Our whole-brain searchlight analysis extended the ROI findings by identifying functional clusters that distinguish between “round” and “spiky” words in the same voxels across participants. One significant cluster was in the left superior temporal gyrus (STG), overlapping with our auditory cortex ROI, an area with a rich history of being linked to complex auditory processing, such as speech perception and comprehension (Buchsbaum et al., 2001; Leff et al., 2009; Ramos Nuñez et al., 2020) and semantic processing (Visser & Lambon Ralph, 2011). With respect to sound symbolism, STG has been shown to exhibit greater univariate activation for round compared to spiky sound-symbolic words when presented concurrently with visual stimuli (Peiffer-Smadja & Cohen, 2019). Without visual stimulation, our results did not indicate differential univariate activity in STG, but instead demonstrate distinct spatially distributed activity patterns, and thus distinct representations, for round and spiky word categories.

The other significant cluster was located in the right intraparietal sulcus (IPS), a region known for its role in visual processing (Swisher et al., 2007), numerical cognition (Duricy et al., 2025), spatial operations (Gillebert et al., 2011; Centanino et al., 2024) and critically involved in the structuring of sensory input across modalities (Cusack, 2005). The latter two functions are particularly relevant for sound symbolism, where auditory input must be mapped onto abstract visuo-spatial properties, and in our study, despite the absence of actual visual stimuli. While previous univariate fMRI studies showed an implication of IPS either when contrasting sound-symbolic versus non-sound-symbolic words (Revill et al., 2014) or when sound-symbolic word-image pairs were incongruent (McCormick et al., 2021), our searchlight results demonstrate distinct neural representations for round and spiky words in right IPS in the absence of visual input.

Together with our ROI results, the identification of STG and IPS clusters could provide a potential functional mechanism on how sound-symbolic speech sounds elicit visual shape representations: Speech sounds are processed and distinguished in primary auditory cortex, left STG and Broca’s area, linked to visual shape representations via visuo-spatial right IPS and eventually fed back to early visual cortex. Thus, the combination of our searchlight and ROI based decoding results provide evidence for the key candidates of brain areas that are involved in the functional pathways giving rise to sound symbolism.

Moreover, our representational similarity analysis revealed a nuanced relationship between behavioral judgments of sound symbolism and their corresponding neural representations. As V1 and left STG exhibited significant correlations between neural pattern similarity and behavioral shape judgments, they may constitute the perceptual-cognitive interface where sound-symbolic mappings are explicitly represented in a format that influences behavior.

The identification of such regions provides insight into where in the processing hierarchy perceptual judgments are formed, moving beyond simply identifying regions that can distinguish between stimulus categories to pinpointing those that display behaviorally meaningful representational geometries. Together with our decoding results, we suggest a functional mechanism in which auditory cortex and Broca’s area analyse acoustic and phonetic features, cross-modal shape associations are activated and represented in right IPS and V1, which in turn leads to corresponding sound symbolic perceptual judgements.

Taken together, the multiple analytical approaches employed in our study advance our understanding of cross-modal representations of sound-symbolic stimuli and provide evidence for a functional neural mechanism underlying the “bouba-kiki” effect. Furthermore, our findings challenge strictly sensory-specific brain organization by revealing the flexible recruitment of purportedly vision-specific cortices for task-relevant cross-modal representations, illustrating how the brain represents information in a more integrated manner than traditionally conceptualized, and converging with evidence that early visual cortex represents several different kinds of abstract auditory information (Vetter et al., 2014; Petro et al., 2017; van den Hurk et al., 2017; Vetter et al., 2020; Mattioni et al., 2020, 2022). The capacity for task-dependent elicitation of shape-specific neural patterns in visual cortices by sound symbolic words could also provide a potential mechanism for the cross-cultural consistency of sound symbolism (Dingemanse et al., 2015; Blasi et al., 2016) .

## Acknowledgements

This work was supported by a research grant from the BIAL Foundation (No. 238/20) to PV and CM, and by a PRIMA grant (PR00P1_185918/1) from the Swiss National Science Foundation to PV. We thank Julia Weber, Fiona Chesnel and Elise Dayer for behavioural piloting of the sound stimuli, Jonas Guggenbühler and Sascha Frühholz for help with sound stimuli recordings, and Bruno Bonet and Damien Marie for scanning support.

## Methods

### Participants

52 healthy adult participants took part in the experiment. Participants were recruited online and received either payment or course credits for their participation. All participants spoke French, had normal or corrected-to-normal vision, normal hearing and were naïve to the “bouba-kiki” effect. They had no neurological or psychiatric history, no claustrophobia, no contradictions for the exposure to the fMRI magnetic field and did not report the use of drugs that might affect neural activity and neurovascular coupling. All research protocols were approved by the Cantonal Ethics Committee of Geneva.

After data quality checks, six participants (four from the shape judgment experiment, two from the semantic judgment experiment) were excluded due to poor model fits (R² < 0.1) in their fMRI general linear model results in response to sounds, regardless of shape condition. Further analysis revealed these participants showed abnormal frequency characteristics in their BOLD signals, with reduced high-to-low frequency power ratios that progressively deteriorated across runs (high/low ratio < 0.02 in final runs compared to > 0.07 in high-quality participants). This suggests a systematic loss of task-relevant neural signal frequencies and accumulation of low-frequency noise, likely explaining their poor model fits. Three of these participants were scanned sequentially, indicating potential scanner issues on that acquisition day. These participants also showed extreme beta values at brain edges, further confirming signal quality issues. While inclusion of these participants did not drastically change the main pattern of results, their exclusion improved the reliability of the analyses by reducing noise. The final sample of 24 participants in the shape judgment experiment (Exp. 1) was composed of 15 females and 9 males, with mean age = 26.5 ± 6.4, and 22 participants in the semantic judgment experiment (Exp. 2) with 18 females and 4 males with mean age = 26.1 ± 7.7.

### Stimuli & task

All sounds were 1 second long two-syllable words and pseudowords, recorded by a male voice with equal stress on both syllables, with balanced frequency of vowels (a, e, i, o, u) and consonants (b, d, g, p, t, k), as reported in Table 1. All sound files were 44.1 kHz, 16-bit WAV files with a mean duration of 1 ± 0.02 seconds and were normalised for amplitude.

Acoustic analysis revealed that word categories nevertheless differed slightly in root mean square amplitude (χ²_3_ = 8.7, *p* = 0.031), despite amplitude normalization. However, the magnitude of this difference (5% between extremes) falls well below the just-noticeable-difference threshold for loudness perception, which is typically 8-12% in amplitude (Jesteadt et al., 1977). Importantly, no significant differences in overall amplitude envelope levels were found (χ²_3_ = 1.9, *p* = 0.59), indicating that overall amplitude levels remained consistent across categories. There were significant spectral differences between word categories (χ²_3_ = 17.5, *p* = 0.0006), with spiky stimuli exhibiting significantly higher spectral centroids (2336 ± 236 Hz) compared to round stimuli (1337 ± 91 Hz), with a gradient across mixed round (1468 ± 233 Hz) and mixed spiky (1953 ± 287 Hz) words. This spectral distinction, trending with the perceived “sharpness” of a sound, resonates with established cross-modal correspondences between auditory frequency and visual shape, where higher frequencies are typically associated with angular shapes (Knoeferle et al., 2017).

The words were pre-validated in a separate behavioural pilot experiment (N = 34) which guided us in the selection of these sound-symbolic words employed in the fMRI experiment (see Results). Shape judgment responses inside the scanner were gathered via an MRI-compatible four-key pad; responses ranged from 1 = round, 2 = mostly round, 3 = mostly spiky to 4 = spiky, or vice versa, counterbalanced across runs. Each run contained 60 trials, 12 per shape condition (each word of Table 1 repeated 2 times) and 12 meaningful words.

Participants completed 4 - 5 runs. A trial comprised 1.4 seconds of stimulus presentation followed by a 4 second inter-trial interval in which participants responded.10% of silent null trials were randomly interleaved.

### MRI acquisition

A 3T Magnetom Prisma scanner (Siemens, Erlangen, Germany) was employed using a 64-channel head coil. Participants underwent anatomical and functional acquisitions during a single session. 3D T1-weighted MPRAGE anatomical volumes were acquired with the following parameters: Repetition Time/Echo Time = 2000/2.49 ms, Inversion Time = 900 ms, Field-of-view = 288 mm^2^, Matrix = 288 × 288 × 192, Slice Thickness = 0.8 mm. All functional MRI series were acquired using a 2D simultaneous multi-slice echo gradient echo planar sequence (2 × 2 mm voxels in-plane; 2 mm slice thickness with no gap; 66 transverse slices, 200 × 200 mm field-of-view; matrix 100 × 100; partial-fourier 7/8; repetition time = 1.32 s; echo time = 30 ms; multiband slice acceleration factor of 3; in-plane acceleration factor of 2; phase encoding direction Anterior-Posterior; flip angle 60°; bandwidth 2174 hz/pixel). For each participant, 4 to 5 runs were gathered, depending on time constraints, with most participants completing 5 runs.

### fMRI preprocessing

Results included in this manuscript come from preprocessing performed using fMRIPrep 22.1.1 (Esteban et al., 2019), which is based on Nipype 1.8.5 (Gorgolewski et al., 2011). A *B_0_* nonuniformity map (or fieldmap) was estimated from the phase-drift map(s) measure with two consecutive GRE (gradient-recalled echo) acquisitions. The T1-weighted (T1w) image was corrected for intensity non-uniformity (INU) (Tustison et al., 2010), distributed with ANTs 2.3.3 (Avants et al., 2011), and used as T1w-reference throughout the workflow. The T1w-reference was then skull-stripped. Brain tissue segmentation of cerebrospinal fluid (CSF), white-matter (WM) and gray-matter (GM) was performed on the brain-extracted T1w (Zhang et al., 2001). Brain surfaces were reconstructed using FreeSurfer 7.2.0,(Dale et al., 1999), and the brain mask estimated previously was refined with a custom variation of the method to reconcile ANTs-derived and FreeSurfer-derived segmentations of the cortical gray-matter of Mindboggle (Klein et al., 2017). Along with the output of images in native space, volume-based spatial normalization to one standard space was performed through nonlinear registration using brain-extracted versions of both T1w reference and the T1w template. The following template was selected for spatial normalization: *ICBM 152 Nonlinear Asymmetrical template version 2009c* (Fonov et al., 2009).

For each of the BOLD runs per participant (across all tasks and sessions), the following preprocessing was performed. First, a reference volume and its skull-stripped version were generated using a custom methodology of fMRIPrep. Head-motion parameters with respect to the BOLD reference (transformation matrices, and six corresponding rotation and translation parameters) were estimated before any spatiotemporal filtering (Jenkinson et al., 2002). The estimated *fieldmap* was then aligned with rigid-registration to the target EPI (echo-planar imaging) reference run. The field coefficients were mapped onto the reference EPI using the transform. The BOLD reference was then co-registered to the T1w reference via boundary-based registration (Greve & Fischl, 2009). Co-registration was configured with six degrees of freedom. Several confounding time-series were calculated based on the preprocessed BOLD: framewise displacement (FD), DVARS and three region-wise global signals. FD was computed using two formulations following Power (Power et al., 2014) (absolute sum of relative motions) and Jenkinson (Jenkinson et al., 2002) (relative root mean square displacement between affines). The three global signals were extracted within the CSF, the WM, and the whole-brain masks. Additionally, a set of physiological regressors were extracted to allow for component-based noise correction (Behzadi et al., 2007).

Principal components were estimated after high-pass filtering the *preprocessed BOLD* time-series (using a discrete cosine filter with 128s cut-off) for the temporal and anatomical variants. Temporal components were then calculated from the top 2% variable voxels within the brain mask. Anatomical components yielded three probabilistic masks (CSF, WM and combined CSF+WM) in native space. The implementation differs from that of Behzadi et al. (2007) in that instead of eroding the masks by 2 pixels on BOLD space, a mask of pixels that likely contain a volume fraction of GM was subtracted from the anatomical components masks. This mask was obtained by dilating a GM mask extracted from the FreeSurfer’s *aseg* segmentation, and it ensured components were not extracted from voxels containing a minimal fraction of GM. Finally, these masks were resampled into BOLD space and binarized by thresholding at 0.99 (as in the original implementation). Components were also calculated separately within the WM and CSF masks. For each decomposition, the *k* components with the largest singular values are retained, such that the retained components’ time series are sufficient to explain 50 percent of variance across the nuisance mask (CSF, WM, combined, or temporal). The remaining components were dropped from consideration. The head-motion estimates calculated in the correction step were also placed within the corresponding confounds file. The confound time series derived from head motion estimates and global signals were expanded with the inclusion of temporal derivatives and quadratic terms for each (Satterthwaite et al., 2013). Frames that exceeded a threshold of 0.5 mm FD or 1.5 standardized DVARS were annotated as motion outliers. Additional nuisance time series are calculated by means of principal components analysis of the signal found within a thin band (*crown*) of voxels around the edge of the brain (Patriat et al., 2017). All resamplings can be performed with a single interpolation step by composing all the pertinent transformations (i.e. head-motion transform matrices, susceptibility distortion correction when available, and co-registrations to anatomical and output spaces). Gridded (volumetric) resamplings were performed using Lanczos interpolation to minimize the smoothing effects of other kernels (Lanczos, 1964).

### Anatomical Regions of Interest

High-resolution ROIs of the early visual cortex were delineated by exploiting the distinctive surface topology of the striate cortex (Benson et al., 2012; Benson & Winawer, 2018) in native space. For all other ROIs except the VWFA, we employed the conventional approach of atlas-based parcellation (Amunts et al., 2020) in standardized space, followed by transformation into native space for subsequent MVPA. For the VWFA, a sphere ROI was drawn at the coordinates [x = -44, y = -58, z = -15] where this functional area has been reported (Dehaene & Cohen, 2011). To ensure anatomical specificity and maintain consistency with the early visual ROIs (which were confined exclusively to gray-matter voxels), we applied gray-matter masking to all other ROIs by generating a binary mask from the FreeSurfer gray-matter probability segmentation, including all voxels with probability values greater than zero. The ROIs varied in size. The EVC, formed by combining V1, V2, and V3, had 3775 ± 481 voxels), with its constituent areas showing comparable sizes (V1: 1237 ± 141; V2: 1312 ± 172; V3: 1225 ± 217 voxels). LO1 (140 ± 29 voxels), LO2 (209 ± 54) and VWFA were significantly smaller (74 ± 18 voxels). The AC, comprising the three subdivisions of the primary auditory region, contained 822 ± 150 voxels in total, with contributions from its component areas (A1: 338 ± 67; A2: 201 ± 45; A3: 282 ± 49 voxels). The PCG was also substantial in size, containing 2544 ± 315 voxels. Finally, Broca’s area was the largest, with 4898 ± 584 voxels on average.

### Analyses

For the behavioral analysis, we extracted and processed participant responses to determine their perceptual judgments of shape characteristics across different stimulus conditions. We then standardized the response coding scheme to ensure consistent interpretation across participants, with lower values corresponding to “round” judgments and higher values to “spiky” judgments on a scale from 1 to 4. We then calculated mean shape ratings for each stimulus condition, allowing us to quantify the perceived “roundness” or “spikiness” of each stimulus category. These data were subsequently analyzed using an analysis of variance (ANOVA) to assess the statistical significance of perceptual differences between stimulus conditions.

For neural response estimation, we implemented GLMsingle (Prince et al., 2022) to generate trial-wise beta maps representing condition-specific neural responses while effectively modeling trial-by-trial variability and mitigating noise.

For the multivariate pattern analysis, we employed a linear support vector machine classification with L2 regularization and default C parameter (C=1.0). Prior to classification, data preprocessing included variance thresholding to remove non-informative voxels and standardization to normalize each feature to zero mean and unit variance (estimated on the training set and applied to both training and test set). A leave-one-run-out cross-validation scheme was employed to evaluate classification performance, effectively testing generalization while respecting the temporal structure of the data, with the final accuracy averaged across folds. For the meaningful versus meaningless classification, we randomly sampled 3 pseudowords from the 4 meaningless shape-associated categories in each run, thereby resulting in balanced classifications of 12 meaningful and 12 meaningless pseudowords. To assess statistical significance, we conducted 1000 iterations of label permutation within each cross-validation fold to generate an empirical null distribution for each participant and ROI. We then calculated chance-subtracted accuracies by subtracting the mean of each participant’s null distribution from their observed classification accuracy. At the group level, we performed Wilcoxon signed-rank tests on these chance-subtracted accuracies, with multiple comparison correction (FDR) applied separately for each classification pair across ROIs.

As an intuitive measure of effect size, we employed a Bayesian approach to quantify the population prevalence (Ince et al., 2021) of this cross-modal effect, with the maximum a posteriori estimate indicating that 31% of the general population in V1, and 63% in A3 would display classification accuracy exceeding the 90th percentile of null distributions derived from 1000 label permutations.

The searchlight analysis employed a spherical kernel with a 3-voxel radius that systematically traversed the gray matter, applying the same preprocessing and classification framework used in our ROI analyses. At each searchlight position, a linear SVM classifier with L2 regularization assessed discriminability between condition pairs using leave-one-run-out cross-validation. The resulting accuracy maps were initially computed in each participant’s native space, then transformed to standard MNI space through a two-step registration process: first resampling to the participant’s T1-weighted anatomical dimensions, then applying the appropriate transformation to align with the MNI template. For group analysis, we standardized the maps by converting to percentage accuracy and subtracting chance level from all non-zero voxels. Group-level statistical inference combined parametric and non-parametric approaches. The parametric analysis employed a second-level general linear model with minimal spatial smoothing (2mm FWHM) to maintain spatial precision. For robust statistical control, we conducted non-parametric permutation testing with 10,000 iterations, implementing a one-sided test focused on above-chance classification accuracy with a stringent initial threshold (p < 0.001) for cluster formation and subsequent cluster-size error rate control.

For the RSA, we constructed neural representational dissimilarity matrices (RDMs) in each ROI by computing pairwise Euclidean distances between beta maps that were averaged per condition, following variance thresholding to remove non-informative voxels. The corresponding behavioral RDMs were derived from participants’ shape judgments, which were standardized across participants to a common scale (1 = round, 4 = spiky). We calculated Kendall’s tau correlation coefficients to quantify the relationship between neural and behavioral RDMs for each participant and region. This non-parametric rank correlation measure assessed whether pairs of conditions that were perceived as more similar behaviorally also produced more similar patterns of neural activity. Group-wise statistical significance was evaluated using Wilcoxon signed-rank tests on the Kendall’s tau correlation coefficients across participants, with correction for multiple comparisons across regions of interest.

## Data & Code

All data and analyses from this study are available for download at https://zenodo.org/records/17448577.

---

**Supplementary Figure S1:**
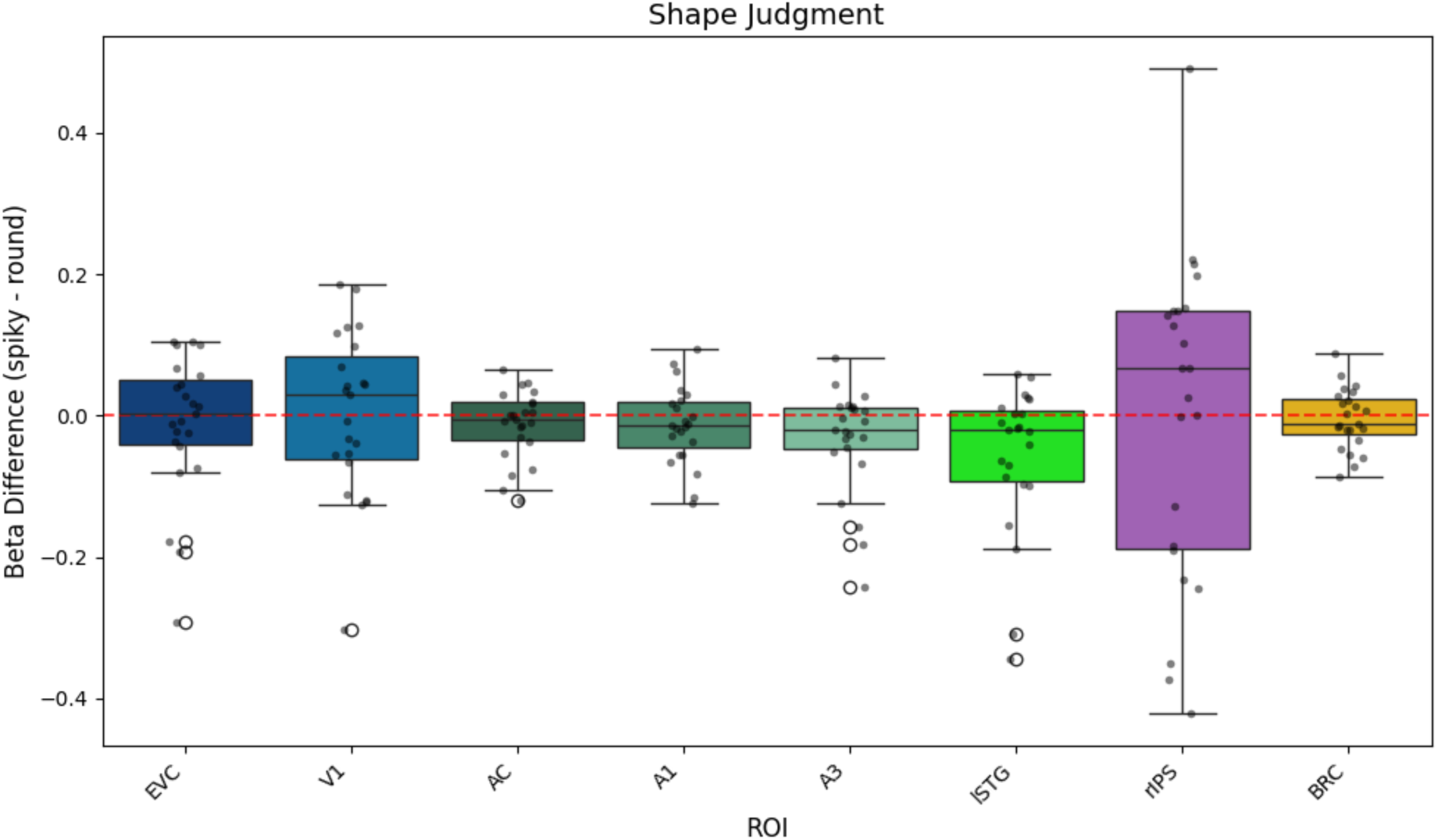
Mean differences in univariate activity levels (beta values) between round and spiky words across relevant ROIs during shape judgment (Experiment 1). In none of these ROIs did mean univariate activity differences depart from 0 (all *p_cor_r* > 0.18).

**Supplementary Figure S2:**
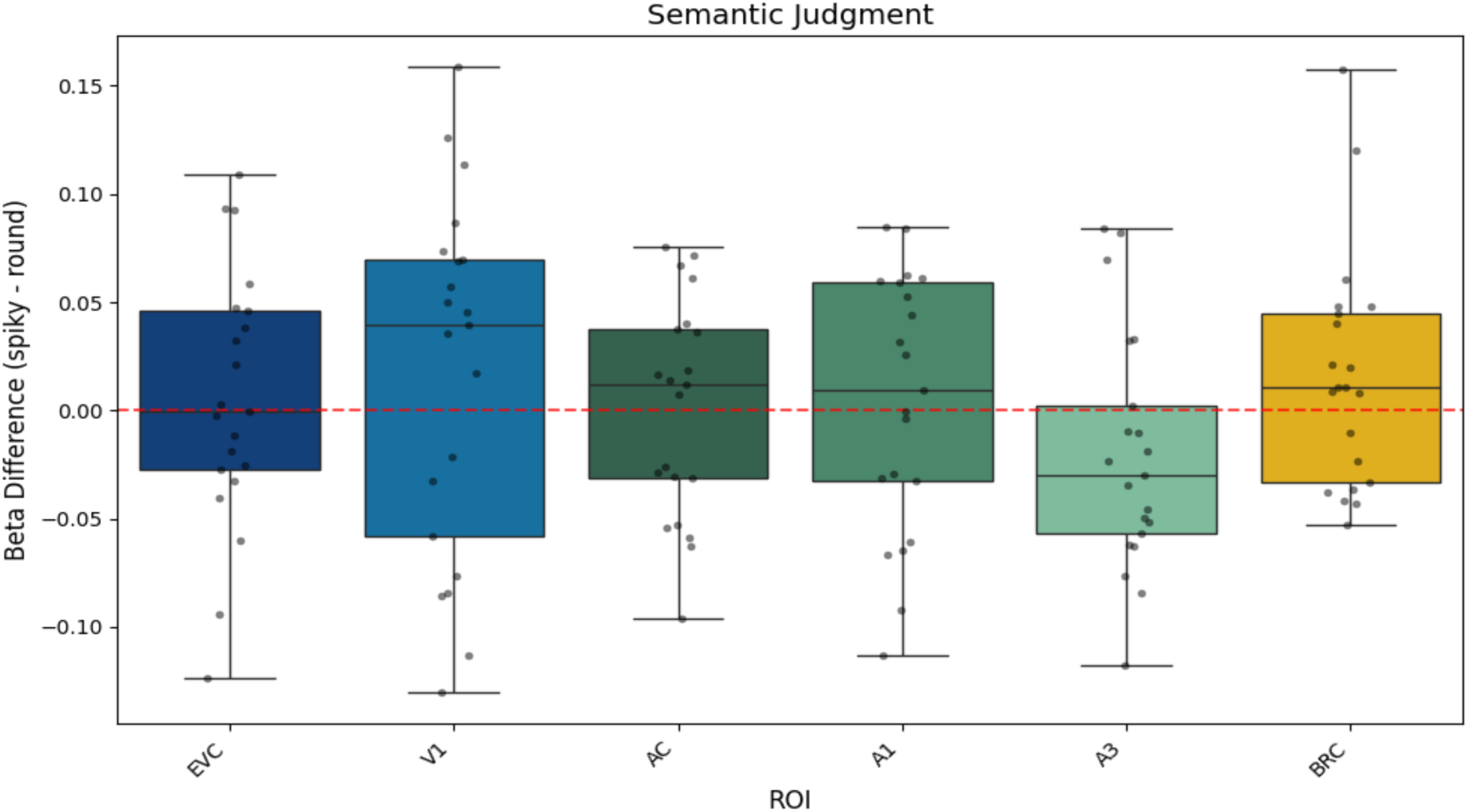
Mean differences in univariate activity levels (beta values) between round and spiky words conditions across relevant ROIs during meaningfulness judgements (Experiment 2). In none of these ROIs did mean activity differences depart from 0 (all *p_corr_* > 0.7).

**Supplementary Figure S3:**
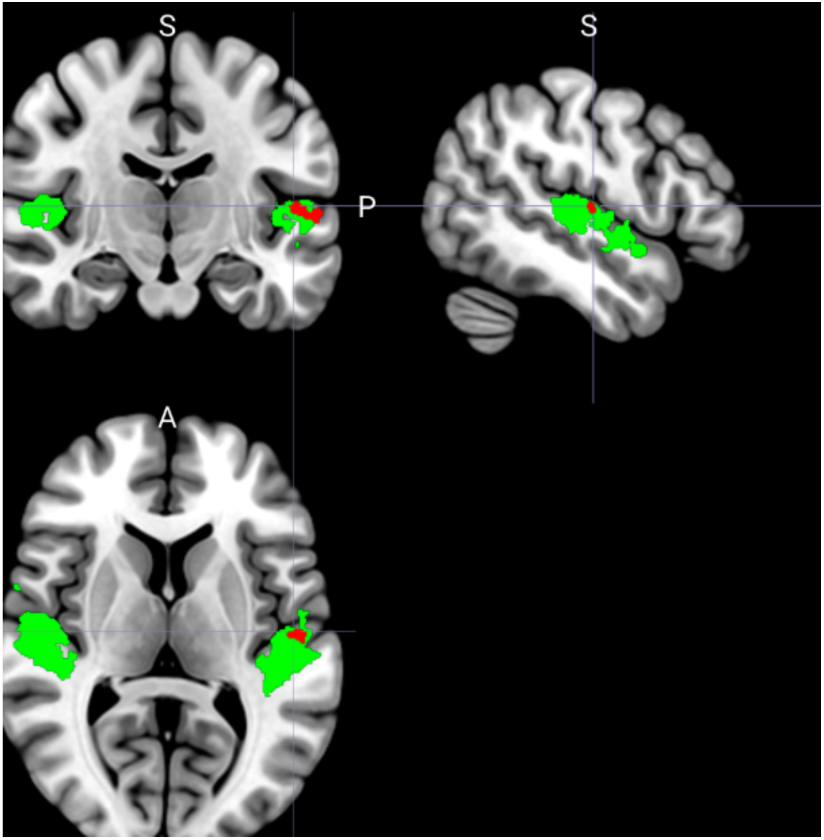
Anatomical overlap of our auditory cortex ROIs (green) with the group-level left STG cluster (red) as identified as distinguishing round versus spiky words in the searchlight analysis.

**Supplementary Figure S4:**
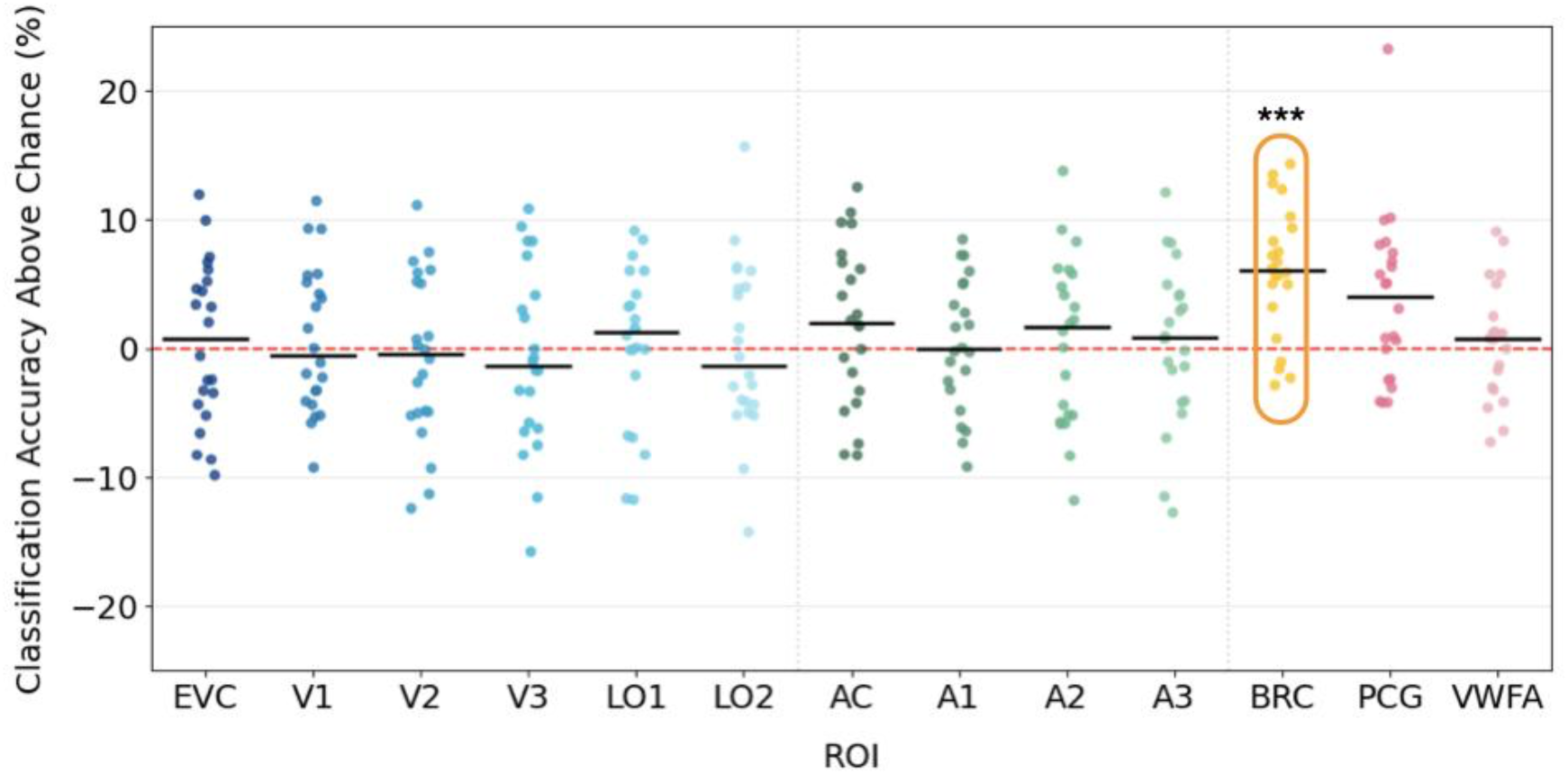
MVPA chance-subtracted accuracies (y axis) for each participant (dots) and ROI (x-axis) for the classification between meaningful and meaningless sounds during meaningfulness judgments (Experiment 2). The red dashed line represents chance level, horizontal black bars represent the group-wise medians and orange contours highlight significant ROIs (\*\*\**p_corr_* <.001) .

**Supplementary Table S1:** Complete MVPA Classification Results - Shape Judgment Task.

| ROI | Classification | p-value | Corrected p-value (FDR) | Significance |
| --- | --- | --- | --- | --- |
| EVC | round_vs_spiky | 0.0020 | 0.0065 | ** |
| EVC | mixed_round_vs_mixed_spiky | 0.6634 | 0.7187 | ns |
| EVC | mixed_round_vs_round | 0.0691 | 0.2727 | ns |
| EVC | mixed_spiky_vs_spiky | 0.8774 | 0.9441 | ns |
| EVC | meaningful_vs_meaningless | 0.3449 | 0.5604 | ns |
| V1 | round_vs_spiky | 0.0020 | 0.0065 | ** |
| V1 | mixed_round_vs_mixed_spiky | 0.8071 | 0.9162 | ns |
| V1 | mixed_round_vs_round | 0.4349 | 0.9162 | ns |
| V1 | mixed_spiky_vs_spiky | 0.8071 | 0.9162 | ns |
| V1 | meaningful_vs_meaningless | 0.8071 | 0.9162 | ns |
| V2 | round_vs_spiky | 0.0173 | 0.0690 | ns |
| V2 | mixed_round_vs_mixed_spiky | 0.1208 | 0.2727 | ns |
| V2 | mixed_round_vs_round | 0.1208 | 0.2727 | ns |
| V2 | mixed_spiky_vs_spiky | 0.6634 | 0.9162 | ns |
| V2 | meaningful_vs_meaningless | 0.0606 | 0.2727 | ns |
| V3 | round_vs_spiky | 0.0173 | 0.0690 | ns |
| V3 | mixed_round_vs_mixed_spiky | 0.3449 | 0.9162 | ns |
| V3 | mixed_round_vs_round | 0.4349 | 0.9162 | ns |
| V3 | mixed_spiky_vs_spiky | 0.6634 | 0.9162 | ns |
| V3 | meaningful_vs_meaningless | 0.8071 | 0.9162 | ns |
| LO1 | round_vs_spiky | 0.0434 | 0.1162 | ns |
| LO1 | mixed_round_vs_mixed_spiky | 0.4349 | 0.5968 | ns |
| LO1 | mixed_round_vs_round | 0.0606 | 0.2727 | ns |
| LO1 | mixed_spiky_vs_spiky | 0.4349 | 0.7187 | ns |
| LO1 | meaningful_vs_meaningless | 0.0606 | 0.2727 | ns |
| LO2 | round_vs_spiky | 0.0173 | 0.0690 | ns |
| LO2 | mixed_round_vs_mixed_spiky | 0.1208 | 0.2727 | ns |
| LO2 | mixed_round_vs_round | 0.8774 | 0.9888 | ns |
| LO2 | mixed_spiky_vs_spiky | 0.8774 | 0.9888 | ns |
| LO2 | meaningful_vs_meaningless | 0.9639 | 0.9888 | ns |
| AC | round_vs_spiky | 0.00001 | 0.00001 | *** |
| AC | mixed_round_vs_mixed_spiky | 0.1554 | 0.4063 | ns |
| AC | mixed_round_vs_round | 0.1554 | 0.4063 | ns |
| AC | mixed_spiky_vs_spiky | 0.3449 | 0.5604 | ns |
| AC | meaningful_vs_meaningless | 0.1554 | 0.4063 | ns |
| A1 | round_vs_spiky | 0.0346 | 0.0934 | ns |
| A1 | mixed_round_vs_mixed_spiky | 0.4349 | 0.7469 | ns |
| A1 | mixed_round_vs_round | 0.0606 | 0.2727 | ns |
| A1 | mixed_spiky_vs_spiky | 0.2056 | 0.4909 | ns |
| A1 | meaningful_vs_meaningless | 0.0606 | 0.2727 | ns |
| A2 | round_vs_spiky | 0.2753 | 0.6325 | ns |
| A2 | mixed_round_vs_mixed_spiky | 0.2056 | 0.4909 | ns |
| A2 | mixed_round_vs_round | 0.2056 | 0.4909 | ns |
| A2 | mixed_spiky_vs_spiky | 0.2056 | 0.4909 | ns |
| A2 | meaningful_vs_meaningless | 0.2056 | 0.4909 | ns |
| A3 | round_vs_spiky | 0.0001 | 0.0001 | *** |
| A3 | mixed_round_vs_mixed_spiky | 0.1554 | 0.4063 | ns |
| A3 | mixed_round_vs_round | 0.1554 | 0.4063 | ns |
| A3 | mixed_spiky_vs_spiky | 0.0692 | 0.2727 | ns |
| A3 | meaningful_vs_meaningless | 0.1554 | 0.4063 | ns |
| PCG | round_vs_spiky | 0.4349 | 0.7469 | ns |
| PCG | mixed_round_vs_mixed_spiky | 0.6634 | 0.9888 | ns |
| PCG | mixed_round_vs_round | 0.8774 | 0.9888 | ns |
| PCG | mixed_spiky_vs_spiky | 0.9639 | 0.9888 | ns |
| PCG | meaningful_vs_meaningless | 0.9639 | 0.9888 | ns |
| BRC | round_vs_spiky | 0.0087 | 0.0249 | * |
| BRC | mixed_round_vs_mixed_spiky | 0.2753 | 0.7469 | ns |
| BRC | mixed_round_vs_round | 0.0606 | 0.2727 | ns |
| BRC | mixed_spiky_vs_spiky | 0.4349 | 0.7469 | ns |
| BRC | meaningful_vs_meaningless | 0.0606 | 0.2727 | ns |
| VWF<br>A | round_vs_spiky | 0.1208 | 0.2843 | ns |
| VWF<br>A | mixed_round_vs_mixed_spiky | 0.4349 | 0.7469 | ns |
| VWF<br>A | mixed_round_vs_round | 0.8774 | 0.9888 | ns |
| VWF<br>A | mixed_spiky_vs_spiky | 0.6634 | 0.9888 | ns |
| VWF<br>A | meaningful_vs_meaningless | 0.9639 | 0.9888 | ns |

**Supplementary Table S2:** Complete MVPA Classification Results - Semantic Judgment Task.

| ROI | Classification | p-value | Corrected p-value (FDR) | Significance |
| --- | --- | --- | --- | --- |
| EVC | round_vs_spiky | 1.0000 | 1.0000 | ns |
| EVC | mixed_round_vs_mixed_spiky | 0.4245 | 0.5518 | ns |
| EVC | mixed_round_vs_round | 0.7262 | 0.8128 | ns |
| EVC | mixed_spiky_vs_spiky | 0.8987 | 0.8987 | ns |
| EVC | meaningful_vs_meaningless | 0.7262 | 0.9193 | ns |
| V1 | round_vs_spiky | 0.5779 | 0.7748 | ns |
| V1 | mixed_round_vs_mixed_spiky | 0.4245 | 0.6632 | ns |
| V1 | mixed_round_vs_round | 0.8987 | 0.9193 | ns |
| V1 | mixed_spiky_vs_spiky | 0.8987 | 0.9193 | ns |
| V1 | meaningful_vs_meaningless | 0.8987 | 0.9193 | ns |
| V2 | round_vs_spiky | 1.0000 | 1.0000 | ns |
| V2 | mixed_round_vs_mixed_spiky | 0.2674 | 0.5518 | ns |
| V2 | mixed_round_vs_round | 0.7262 | 0.8128 | ns |
| V2 | mixed_spiky_vs_spiky | 0.8987 | 0.9193 | ns |
| V2 | meaningful_vs_meaningless | 0.8987 | 0.9193 | ns |
| V3 | round_vs_spiky | 0.5779 | 0.7748 | ns |
| V3 | mixed_round_vs_mixed_spiky | 0.5779 | 0.7748 | ns |
| V3 | mixed_round_vs_round | 0.7262 | 0.8128 | ns |
| V3 | mixed_spiky_vs_spiky | 0.5779 | 0.7748 | ns |
| V3 | meaningful_vs_meaningless | 0.8987 | 0.9193 | ns |
| LO1 | round_vs_spiky | 0.5779 | 0.7748 | ns |
| LO1 | mixed_round_vs_mixed_spiky | 0.4245 | 0.6632 | ns |
| LO1 | mixed_round_vs_round | 0.2674 | 0.5518 | ns |
| LO1 | mixed_spiky_vs_spiky | 0.7262 | 0.8128 | ns |
| LO1 | meaningful_vs_meaningless | 0.8987 | 0.9193 | ns |
| LO2 | round_vs_spiky | 0.5779 | 0.7748 | ns |
| LO2 | mixed_round_vs_mixed_spiky | 0.4245 | 0.6632 | ns |
| LO2 | mixed_round_vs_round | 0.8987 | 0.9746 | ns |
| LO2 | mixed_spiky_vs_spiky | 0.8987 | 0.9746 | ns |
| LO2 | meaningful_vs_meaningless | 0.8987 | 0.9746 | ns |
| VWF<br>A | round_vs_spiky | 0.5779 | 0.7748 | ns |
| VWF<br>A | mixed_round_vs_mixed_spiky | 0.4245 | 0.6632 | ns |
| VWF<br>A | mixed_round_vs_round | 0.8987 | 0.9193 | ns |
| VWF<br>A | mixed_spiky_vs_spiky | 0.7262 | 0.8128 | ns |
| VWF<br>A | meaningful_vs_meaningless | 0.8987 | 0.9193 | ns |
| AC | round_vs_spiky | 0.0001 | 0.0001 | *** |
| AC | mixed_round_vs_mixed_spiky | 0.1327 | 0.4245 | ns |
| AC | mixed_round_vs_round | 0.0152 | 0.0690 | ns |
| AC | mixed_spiky_vs_spiky | 0.2674 | 0.5518 | ns |
| AC | meaningful_vs_meaningless | 0.8987 | 0.9193 | ns |
| A1 | round_vs_spiky | 0.0001 | 0.0003 | *** |
| A1 | mixed_round_vs_mixed_spiky | 0.0152 | 0.0690 | ns |
| A1 | mixed_round_vs_round | 0.0690 | 0.2674 | ns |
| A1 | mixed_spiky_vs_spiky | 0.0357 | 0.1327 | ns |
| A1 | meaningful_vs_meaningless | 0.8987 | 0.9193 | ns |
| A2 | round_vs_spiky | 0.0152 | 0.0542 | ns |
| A2 | mixed_round_vs_mixed_spiky | 0.0357 | 0.1327 | ns |
| A2 | mixed_round_vs_round | 0.0357 | 0.1327 | ns |
| A2 | mixed_spiky_vs_spiky | 0.1327 | 0.4245 | ns |
| A2 | meaningful_vs_meaningless | 0.8987 | 0.9193 | ns |
| A3 | round_vs_spiky | 0.0044 | 0.0159 | * |
| A3 | mixed_round_vs_mixed_spiky | 0.0152 | 0.0690 | ns |
| A3 | mixed_round_vs_round | 0.0152 | 0.0690 | ns |
| A3 | mixed_spiky_vs_spiky | 0.0152 | 0.0690 | ns |
| A3 | meaningful_vs_meaningless | 0.8987 | 0.9193 | ns |
| PCG | round_vs_spiky | 0.5779 | 0.7748 | ns |
| PCG | mixed_round_vs_mixed_spiky | 0.2674 | 0.5518 | ns |
| PCG | mixed_round_vs_round | 0.4245 | 0.6632 | ns |
| PCG | mixed_spiky_vs_spiky | 0.4245 | 0.6632 | ns |
| PCG | meaningful_vs_meaningless | 0.0152 | 0.0671 | ns |
| BRC | round_vs_spiky | 0.0152 | 0.0542 | ns |
| BRC | mixed_round_vs_mixed_spiky | 0.2674 | 0.5518 | ns |
| BRC | mixed_round_vs_round | 0.0357 | 0.1327 | ns |
| BRC | mixed_spiky_vs_spiky | 0.1327 | 0.4245 | ns |
| BRC | meaningful_vs_meaningless | 0.0001 | 0.0007 | *** |

